# Transcriptional Heterogeneity Reveals a Synaptic Gene Program in Developing and Adult Human Oligodendrocyte Precursor Cells

**DOI:** 10.64898/2026.01.01.697017

**Authors:** Aliya R. Grinberg, Li Liu, Irva Patel, Brinda Guntur, Ian A. Glass, Birth Defects Research Laboratory, Kosaku Shinoda, Hiroko Nobuta

**Author notes:** Corresponding authors: Kosaku Shinoda; Hiroko Nobuta.

## Abstract

Human oligodendrocyte precursor cells (OPCs) arise early in gestation and expand broadly during cortical development, yet the extent of their heterogeneity remains poorly defined. Here, we isolated >2,300 highly pure OPCs from post-conceptional week (PCW) 17 human cortex using an optimized PDGFRα-based immunopanning and performed single-cell RNA sequencing. Unsupervised clustering revealed four transcriptionally distinct embryonic OPC subsets, including a previously unrecognized population that expressed genes linked to synaptic development, synaptic signaling, and neuromodulation. This subset - designated embryonic synaptic OPCs (eSyn-OPCs) - comprised approximately 28.5% of all embryonic OPCs in the cortex and was characterized by robust expression of synapse-associated secreted factors (*THBS2, WNT5A, WNT7A, PLAT, ACHE*) and multiple neurotransmitter receptor subunits. Histological analyses across PCW 12–22 demonstrated that eSyn-OPCs first appear around PCW 15 and are enriched in proliferative germinal zones. Spatial transcriptomics confirmed their localization near neural stem and progenitor cells, suggesting proximal neuron-OPC communication during early cortical assembly. Purified eSyn-OPCs differentiated into mature oligodendrocytes *in vitro*, confirming their oligodendrocyte lineage identity. Reanalysis of adult human single-nucleus RNA-seq datasets uncovered a transcriptionally analogous OPC subset (adult synaptic OPCs, aSyn-OPCs), though with reduced representation of structural synaptic genes and neurotransmitter receptor diversity compared to eSyn-OPCs. Together, these results identify a synaptically specialized OPC population in both developing and adult human cortex and reveal that eSyn-OPCs possess unexpectedly rich synaptic signaling machinery. These findings suggest that human OPCs may participate directly in neuron-glia communication during early cortical development and raise the possibility of developmental stage-specific roles for eSyn-OPCs in shaping neural circuit formation.

## INTRODUCTION

Accumulating evidence suggests that human oligodendrocyte precursor cells (OPCs) are generated as early as post-conceptional week (PCW) 6 in the hindbrain [1], and PCW 8 in the ventral forebrain [2]. They undergo dramatic expansion in the 2^nd^ trimester and become widely distributed across cortical layers [3, 4]. Differentiated OPCs contribute to myelination of neuronal axons beginning in late 2^nd^ trimester [5, 6] and continues into young adulthood [7, 8]. OPCs remain in human brains throughout life [9, 10], and they are thought to participate in remyelination after injury [11–14]. Historically, a largely homogenous OPC population was thought to constitute myelin, but recent studies demonstrated that in mouse, transcriptionally and functionally distinct subtypes of OPCs contribute to myelin [15, 16]. However, it is unclear whether human OPC population is homogeneous or comprised of heterogenous subtypes.

We used an optimized immunopanning method to collect >2,300 highly pure human OPCs from post-conceptual week (PCW) 17 brain and analyzed single-cell transcriptome. We identified distinct clusters of OPCs based on transcriptomic and signaling characteristics. One cluster, we termed syn-OPCs, showed unique genes related to synaptic development and signaling. Histological analyses of cortical tissues across the 2^nd^ trimester demonstrated that syn-OPCs appear in germinal zones (subventricular, inner ventricular, and outer ventricular zones) at around PCW15. Spatial transcriptomics analyses of cortical tissues showed that syn-OPCs are potentially interacting with young neurons in these regions. Using a publicly available dataset of adult human brain cells [17], we identified a similar OPC cluster with synaptic signaling roles. However, a detailed examination revealed distinct molecular characteristics compared to embryonic syn-OPCs highlighting the abundant expression of multiple neurotransmitter receptor genes in embryonic syn-OPCs. These observations suggest that a subcluster of human OPCs have the potential to receive multiple neurotransmitters released by neurons during active neuronal differentiation.

## RESULTS

### Transcriptional heterogeneity of human embryonic OPCs

Due to their fragility, isolating large numbers of OPCs from intact human tissues has long been challenging [3]. We optimized an immunopanning protocol using PDGFRα (CD140a) antibody to improve the yield of isolated human OPCs (see Methods). With this approach, we obtained highly pure (>90%) and viable (>92%) OPCs from a post-conceptional week (PCW)17 cortical specimen [18]. The isolated cells were subjected to droplet-based single-cell RNA sequencing (scRNA-seq), and only those that passed rigorous quality controls criteria were analyzed.

A total of 2,304 high-quality cells were obtained (**Fig. 1A**). These cells expressed *PDGFRA* and *OLIG2* abundantly (**Fig. 1B**) but lacked mature oligodendrocyte markers such as *MOG* and *MAG* (**Fig. 1C**), confirming the OPC identity. Unsupervised clustering based on transcriptomic similarity revealed four well-defined OPC clusters (**Fig. 1D**). As shown, each cluster exhibited a unique and largely non-overlapping gene expression signature. For convenience, we designated these four clusters as <u>e</u>mbryonic (e) Clusters 1 – 4 (**Fig. 1A, D-E**). Gene Ontology (GO) enrichment analysis of differentially expressed genes (DEGs) showed that eCluster 1 was associated with brain development–related pathways (**Fig. 1E**). Consistent with the extensive cellular expansion known to take place in the embryonic cortex [3, 19–21], we identified multiple proliferative OPC clusters. As shown in **Fig. 1F**, eCluster 2 highly expressed cell cycle and mitosis genes indicative of active proliferation. Although overlapped to some extent to eCluster 2 in cell cycle genes, eCluster 3 specifically expressed genes related to chromosome separation and centromere assembly (**Fig. 1G**), suggesting the cells are in G1-to-S phase. These eClusters 1 - 3 represented OPC subsets consistent with expected developmental stages of embryonic OPC generation.

**Figure 1.**
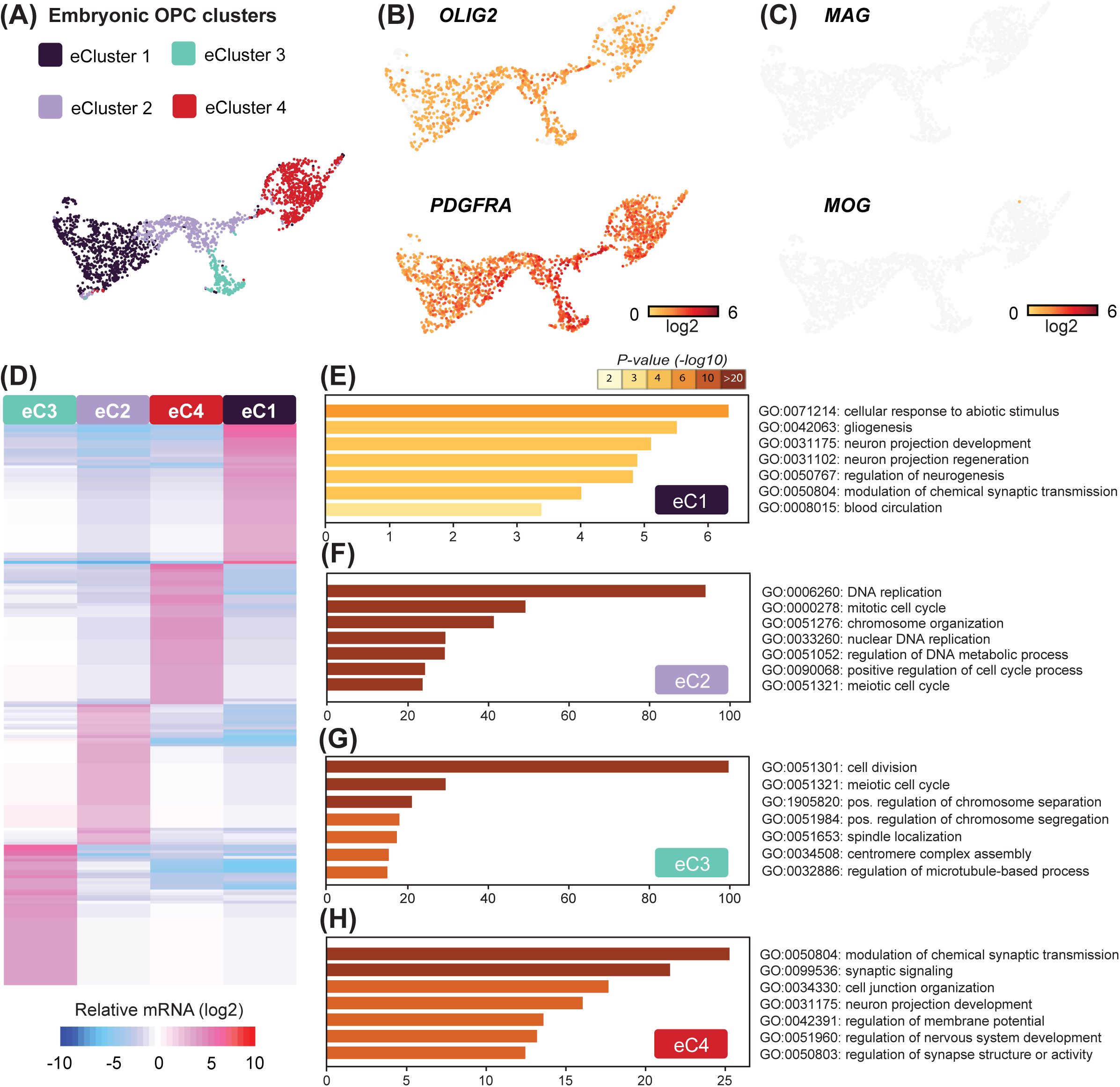
Characterization of OPC heterogeneity in PCW 17 human cortex. **(A)** Human embryonic (PCW 17) OPCs were isolated based on PDGFRα immunoreactivity (see Methods) and subjected to scRNA-seq. Graph clustering based on transcriptomic similarity revealed four OPC clusters (<u>e</u>mbryonic clusters 1-4) in UMAP space. **(B)** Cells showed robust expression of *OLIG2* and *PDGFRα*, confirming OPC identity. **(C)** Mature oligodendrocyte markers *MAG* or *MOG* were low. **(D)** Heatmap representation of a unique and largely non-overlapping gene expression signatures for each OPC cluster. **(E-H)** Top seven pathways identified by Gene Ontology (GO) analysis of differentially expressed (*P* < 0.01) genes in each OPC cluster.

In contrast, we identified an unanticipated cluster, eCluster 4, which expressed genes involved in synaptic transmission, synaptic signaling, and the regulation of synapse structure or activity (**Fig. 1H**, discussed below). At the individual gene level, eCluster 4 expressed *THBS2* (Thrombospondin 2) [22], *PLAT* (Plasminogen activator tissue type) [23]*, WNT5A* (Wnt Family Member 5A) [24, 25]*, WNT7A* (Wnt Family Member 7A) [26, 27], and *ACHE* (Acetylcholinesterase) [28, 29] (**Suppl Fig. 1A-B**), all of which encode for well-established regulators of synaptic development and neuromodulatory secreted factors. Based on this distinctive expression pattern, we designated eCluster 4 as the <u>e</u>mbryonic <u>syn</u>aptic <u>OPC</u> or “eSyn-OPC”. eSyn-OPC comprised approximately 28.5% of all OPCs detected in the PCW 17 cortical tissue (**Fig. 1A**). Importantly, the presence of eSyn-OPCs was validated in independent cortical samples across 2^nd^ trimester (see **Fig. 4C**).

### Similarities and divergence of embryonic vs adult OPCs

Are eSyn-OPCs a transient population restricted to the developing brain, or do they also exist in the adult brain? To address this question, we analyzed a publicly available single-nucleus RNA-seq dataset derived from the cortex of a 29-year-old male who had died suddenly from pulmonary embolism [17]. This dataset encompassed all major brain cell classes (superclusters), including excitatory and inhibitory neurons, microglia, astrocytes, endothelial cells, and fibroblasts. We confidently identified OPCs within the superclusters by the co-expression of *PDGFRA* and *OLIG2*, the definitive molecular markers of OPCs (**Fig. 2A**). This OPC supercluster was clearly delineated from mature oligodendrocytes expressing canonical markers such as *MAG*, *MOG*, and *MBP*. The identified OPC supercluster was then re-clustered based on DEGs. Similar to the embryonic OPCs, adult OPCs segregated cleanly into four transcriptionally distinct clusters by unsupervised clustering (**Fig. 2B**). For clarity, we designated these as <u>a</u>dult (a) Clusters 1 - 4.

**Figure 2.**
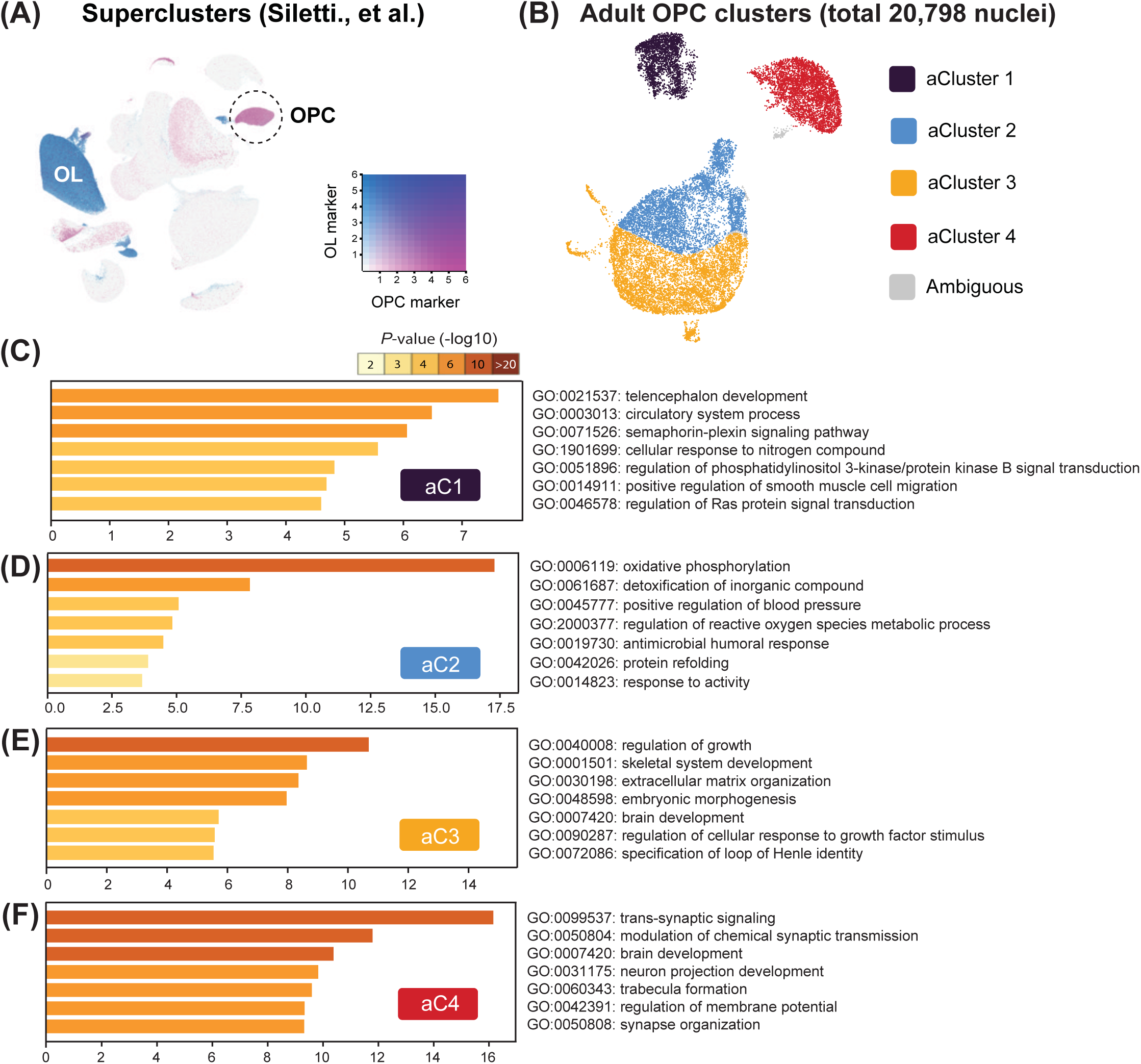
Evidence of Syn-OPCs in the adult human brain. **(A)** Identification of OPCs in the UMAP plot of brain cells obtained from a healthy 29-year-old male (from Siletti et al.). OPCs were defined by co-expression of marker genes *PDGFRA* and *OLIG2*, and mature oligodendrocytes were defined by co-expression of marker genes *MAG*, *MOG*, and *MBP*. **(B)** Re-clustering of the OPC cluster found in (A) based on differentially expressed genes identified four OPC clusters (<u>a</u>dult clusters 1-4). **(C-F)** Top seven pathways identified by GO analysis of differentially expressed genes in each OPC cluster from (B). aCluster 4 was identified as aSyn-OPCs.

aCluster 1 exhibited general OPC-associated biological pathways (**Fig. 2C**), resembling eCluster 1 from the embryonic dataset (**Fig. 1E**). Contrary to embryonic OPCs (eCluster 2 and 3, **Fig. 1F-G**), no proliferative clusters were detected among adult OPCs. Instead, adult OPCs contained unique, non-proliferative subsets characterized by distinct metabolic and growth-related transcriptional programs (**Fig. 2D-E**); aCluster 2 showed enrichment for oxidative phosphorylation and related metabolic pathways, suggesting homeostatic maintenance of adult OPCs (**Fig. 2D**), whereas aCluster 3 was enriched for genes involved in extracellular matrix organization, growth regulation, and brain development, implying a potential trajectory toward differentiation or maturation (**Fig. 2E**). Consistent with the embryonic data, aCluster 4 exhibited significant enrichment in synapse-associated pathways, including trans-synaptic signaling and modulation of chemical synaptic transmission (**Fig. 2F**). Based on this observation, we designated this subset as <u>a</u>dult <u>syn</u>aptic OPCs (aSyn-OPCs).

An important question is whether <u>a</u>Syn-OPCs and <u>e</u>Syn-OPCs represent independent populations that coincidentally share synaptic gene enrichment, or they are biologically related lineages. To assess this, we performed a Monte Carlo simulation (*N* = 10,000) [30] to construct an empirical null distribution of gene-set overlaps. Specifically, we compared the set of 800 marker genes (*P*<0.01) defining eSyn-OPCs (identified in **Fig. 1**) with randomly sampled gene sets of equal size drawn from all genes expressed in the human cortex. The mean overlaps between random gene sets and the eSyn-OPC markers was 3.62% (SD = 0.64). In contrast, 5.59% of genes enriched in aSyn-OPCs overlapped with the eSyn-OPC markers. Although this may appear modest, the Monte Carlo-derived null distribution (**Fig. 3A**) indicates that this value lies more than three standard deviations above the mean, corresponding to the top 0.1% of the null distribution. As a comparison, other adult OPC clusters showed substantially lower overlap rates with eSyn-OPC genes: aCluster 1 exhibited an overlap of 3.81% (*P* = 0.389), essentially matching the null expectation, while aCluster 3 (2.16% overlap) and aCluster 2 (1.97% overlap) fell within the null distribution and are clearly lower than that observed for aSyn-OPCs. To further validate the presence of Syn-OPCs in the adult human brain, we analyzed an additional dataset from another male individual who died at 42 years old using the same bioinformatic framework (**Suppl. Fig. 2**). After re-clustering of the OPC supercluster (**Suppl. Fig. 2A-B**), we confirmed a similar aSyn-OPC cluster (**Suppl. Fig. 2F**). Above findings were replicated in an independent snRNA-seq dataset published by a different group [31]; across all six adult individuals examined (ages 34 - 68 years), we consistently detected a subset of OPCs expressing high levels of synapse-associated genes, spatially segregated from the main OPC cluster in the UMAP plot (**Suppl. Fig. 3A-C**). Thus, across eight adult human brains, we consistently identified a subcluster resembling the eSyn-OPCs, supporting the notion that eSyn-OPC-like cells persist in adulthood.

**Figure 3.**
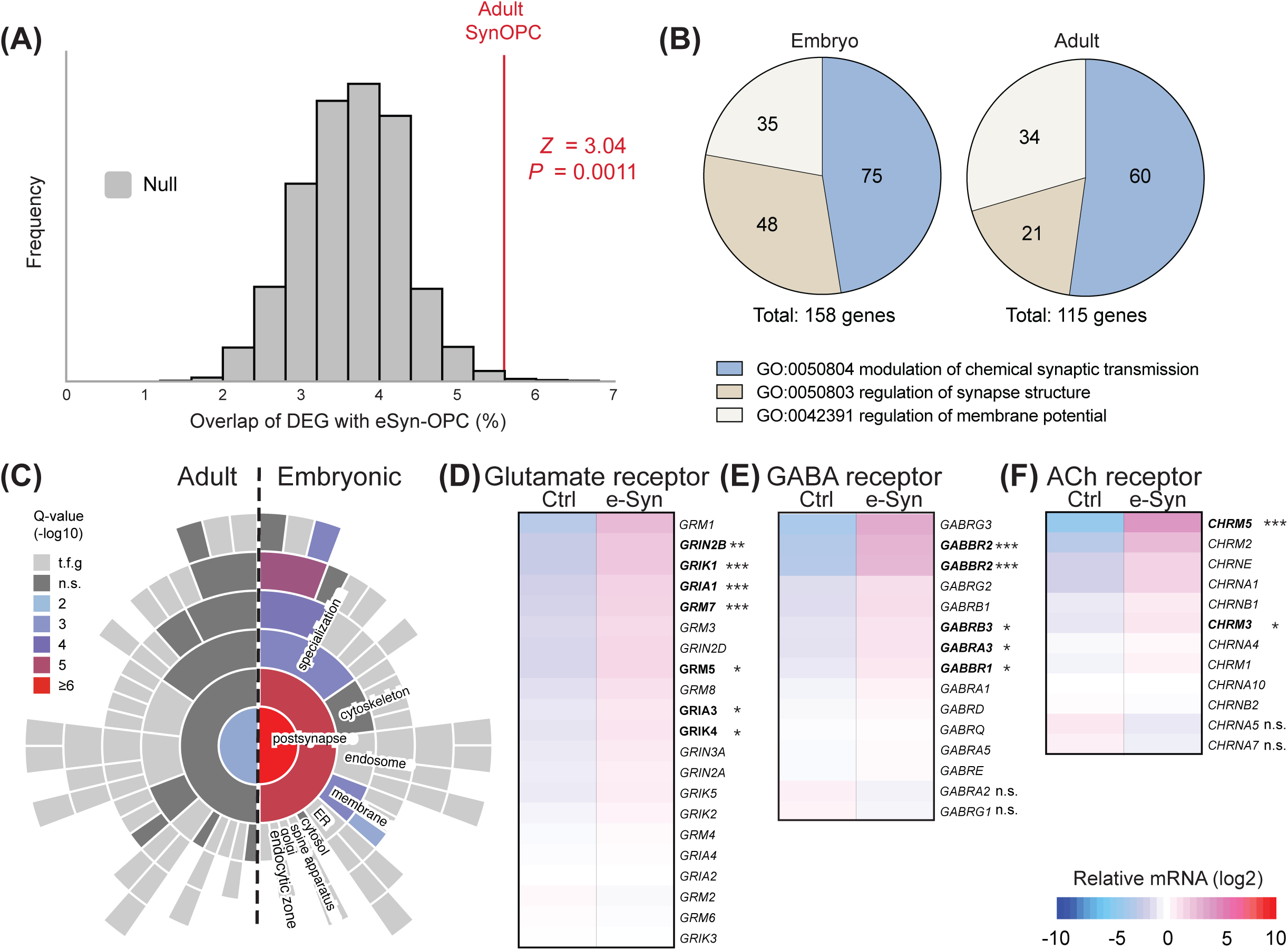
Molecular evidence of putative OPC-neuron communication. **(A)** A simulated null distribution of gene-set overlaps between eSyn-OPCs and randomly sampled gene sets of the same size expressed in the brain (*n* = 10,000). Adult Syn-OPCs (aSyn-OPCs) showed a significant overlap (z = 3.04, p = 0.0011) with eSyn-OPCs, indicating genetic similarity between the two populations. **(B)** Among synapse-related genes, eSyn-OPCs showed a greater abundance of genes regulating synapse structure compared to aSyn-OPCs. **(C)** SynGO, a synaptic gene analysis tool, showed that eSyn-OPCs expressed more genes encoding postsynaptic proteins compared to aSyn-OPCs. **(D-F)** The mRNA levels of postsynaptic neurotransmitter receptor subunit genes for glutamate, GABA, and acetylcholine in eSyn-OPCs compared with other embryonic OPC clusters.

A deeper comparative investigation, however, also revealed a clear difference between eSyn-OPCs and aSyn-OPCs; whereas genes associated with chemical synaptic transmission and membrane potential regulation were similarly enriched in both eSyn-OPCs and aSyn-OPCs (**Fig. 1H & Fig. 2F**), genes regulating synapse structure were abundant by more than two-fold in eSyn-OPCs (**Fig. 3B**). This distinction suggests that eSyn-OPCs may play developmentally specific roles in shaping neuronal connectivity. To dissect this structural program, we utilized SynGO [32] to map eSyn-OPC markers to curated synaptic ontologies. This analysis revealed a marked enrichment of genes encoding postsynaptic proteins in eSyn-OPCs relative to other OPC subclusters (**Fig. 3C**). Specifically, we identified significant upregulation of glutamate (*GRIN2B, GRIK1, GRIA1, GRM7*), GABA (*GABBR2, GABRB3, GABRA3*), and acetylcholine (*CHRM5, CHRM3*) receptor subunit genes compared to conventional OPCs (**Fig. 3D-F**). These findings identify the broad expression of neurotransmitter receptor machinery as a unique characteristic of eSyn-OPCs.

### Temporal and regional specificity of eSyn-OPCs

To define the spatiotemporal localization of eSyn-OPCs, we screened commercially available antibodies targeting the top 15 eSyn-OPC marker genes. This yielded five high-fidelity candidates (EDIL3, GRB14, HAPLN1, IL1RAP, MT1E) that exhibited specific immunoreactivity with minimal background (**Suppl Fig. 4B**). Using these antibodies in combination with a known pan-OPC marker PDGFRα, we validated the utility of these antibodies for detecting eSyn-OPCs by immunohistochemistry. Analysis of adjacent sections from a PCW 22 specimen revealed high concordance across the five antibodies. While minor variability was observed, the proportion of detected eSyn-OPCs (PDGFRα^+^/Marker^+^) was consistent (mean detection of 62.2% of all PDGFRα^+^ OPCs ± 1.99% SEM within outer subventricular zone, **Suppl Fig. 4A-B**). We therefore concluded that these antibodies can be used interchangeably for identifying eSyn-OPCs. To determine the developmental trajectory of eSyn-OPCs, we performed histological assessments using HAPLN1 antibody (**Fig. 4A-C**). Evaluation of cortical tissues spanning PCW 12 to 22 indicated a specific developmental window: eSyn-OPCs were consistently identified between PCW 15 and 22 (**Fig. 4C**). At PCW 12, eSyn-OPC detection was negligible despite the presence of HAPLN1^-^/PDGFRα^+^ OPCs, indicating a later developmental onset for eSyn-OPCs (see Discussion for limitations). These findings were corroborated using an independent marker, GRB14 (data not shown). In tissues spanning PCW 15 to 22 (*n*=3, both sexes), eSyn-OPC distribution varied by age: earlier stages showed enrichment in the ventricular zone (VZ) and inner subventricular zone (iSVZ), whereas older tissues showed localization primarily in the outer subventricular zone (oSVZ) and intermediate zone (IZ) (**Fig. 4A-C**). This pattern is consistent with their putative origin in the VZ followed by migration toward the oSVZ. In striking contrast to conventional OPCs (HAPLN1^-^/PDGFRα^+^), which were distributed throughout the cortical layers, eSyn-OPCs were virtually absent from the upper layers, including the cortical plate and marginal zone. Therefore, we conclude that eSyn-OPCs are predominantly confined to the ventral proliferative zones (VZ, iSVZ, and oSVZ), with the upper layers remaining largely devoid of eSyn-OPCs. (**Fig. 4B-C**). Morphologically, eSyn-OPCs exhibited a simplified architecture with few processes (0–2), whereas HAPLN1^-^ OPCs displayed significantly more elaborate process branching (**Fig. 4B**).

**Figure 4.**
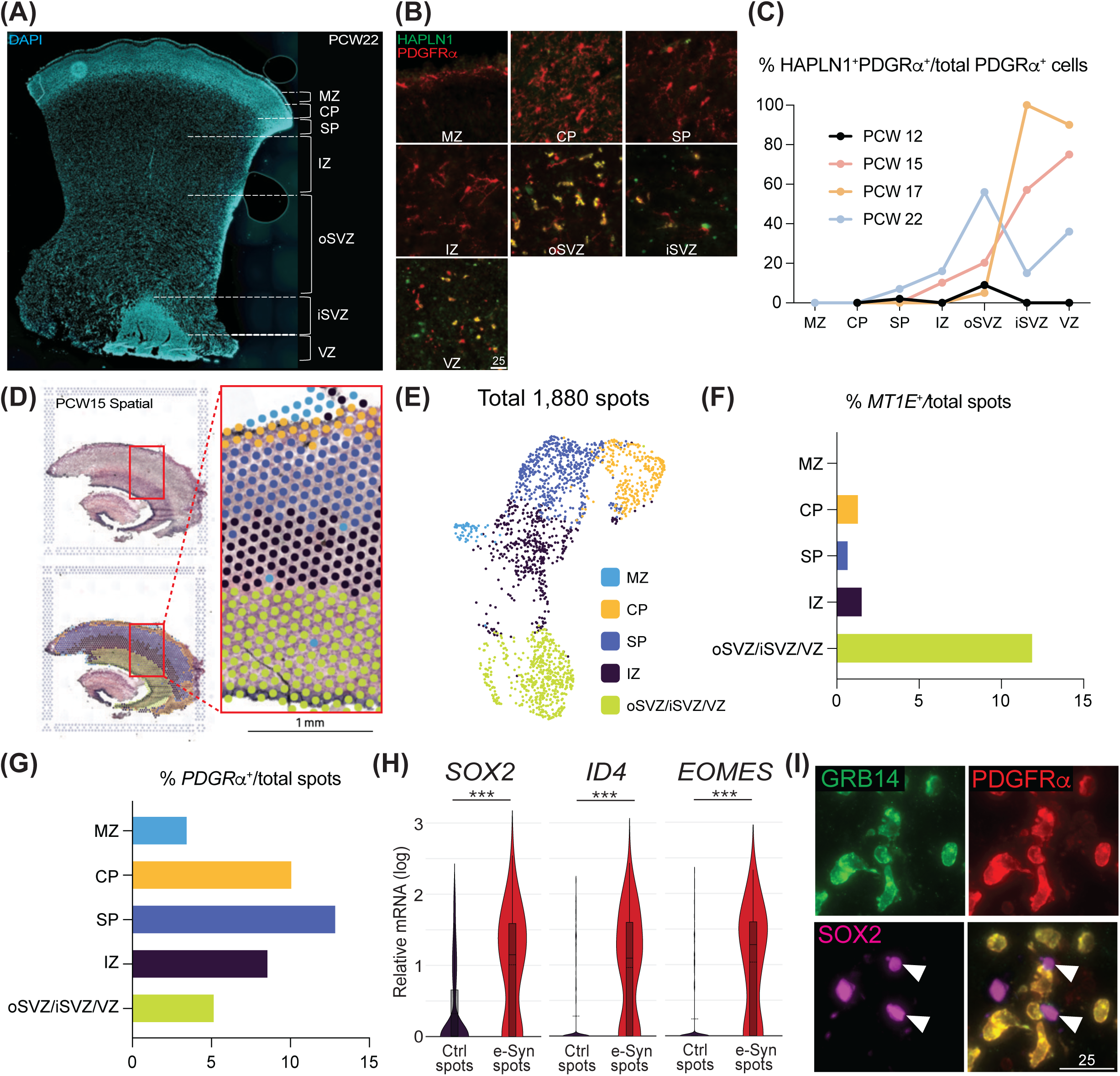
Spatial distribution of eSyn-OPCs and their adjacency to developing neurons. **(A)** A representative cortical section with DAPI staining from a PCW 22 brain. **(B)** Immunohistochemistry for eSyn-OPCs using the eSyn-OPC marker HAPLN1 and colocalizing with PDGFRα. Scale bar unit = um. **(C)** Quantification of HAPLN1^+^/PDGFRα^+^ eSyn-OPCs revealed that the appearance of eSyn-OPCs occurs after PCW 12, and the majority of eSyn-OPCs are found in layers IZ, oSVZ, iSVZ, and VZ. **(D)** Visium spatial transcriptomics from a PCW 15 sample. **(E)** K-means clustering of 1,880 spots from (D) identified five distinct layers in UMAP space. **(F)** Quantification of *MT1E*^+^ eSyn-OPC spots in the five cortical layers found in (E). **(G)** Quantification of *PDGFRα*^+^ OPC spots in the five cortical layers found in (E). **(H)** The mRNA levels of neural progenitor (*EOMES*) and neural stem cell (*SOX2, ID4*) markers in the *MT1E*^+^ spots (containing eSyn-OPCs) compared to *PDGFR*α^+^/*MT1E*^-^ spots (containing OPCs but not eSyn-OPCs). **(I)** Immunohistochemistry confirmed proximity of eSyn-OPCs and SOX2^+^ neural stem cells in PCW 22 cortical tissue. Scale bar unit = um.

We validated these spatiotemporal findings using Visium spatial transcriptomics on fresh-frozen cortical tissues from PCW 8, 15, and 19. *MT1E* was selected as the surrogate marker for eSyn-OPCs because it was more robustly detected than *HAPLN1* in the Visium dataset. This substitution is justified by the fact that MT1E and HAPLN1 exhibit highly concordant expression profiles for eSyn-OPCs in immunohistochemical validation (**Suppl Fig. 4A-B**). Corroborating our scRNA-seq (**Suppl. Fig. 4C**) and histological analyses (**Fig. 4B–C**), the eSyn-OPC marker *MT1E* was significantly enriched in spatial transcriptomic spots from PCW 15 (**Fig. 4D–F**) and 19 (**Suppl Fig. 4E–F**). Conversely, the PCW 8 tissue displayed only negligible *MT1E* expression (8 spot; 0.7%) despite the robust detection of PDGFRα^+^ spots (**Suppl Fig. 4D**). This scarcity aligns with the histological absence of eSyn-OPCs at PCW 12 (**Fig. 4C**), confirming that eSyn-OPCs are virtually absent during the first trimester. Unsupervised clustering of 1,880 Visium spots from the PCW 15 sample identified five distinct transcriptional clusters (**Fig. 4E**). When spatially projected onto the original hematoxylin and eosin section, these clusters faithfully recapitulated the laminar architecture of the developing cortex (**Fig. 4D**). Within this spatial framework, eSyn-OPC spots comprised approximately 12% of the deep germinal zones (oSVZ/iSVZ/VZ in **Fig. 4E-F**), with their prevalence progressively diminishing dorsally and reaching zero spot in the MZ (**Fig. 4E-F**). Conventional OPC spots distributed across cortical layers (**Fig. 4G**). Similarly, PCW 19 sample showed accumulation of eSyn-OPC spots in lower cortical layers (**Suppl Fig. 4E–F**). Taken together, these observations identify eSyn-OPCs as a prominent subcluster predominantly localized to the ventral proliferative zones during the second trimester of human cortical development.

While immunohistochemistry provided high-resolution localization of eSyn-OPCs (**Fig. 4A–C**), identifying their specific interaction partners remains challenging. We therefore analyzed the Visium dataset to identify the cell types residing in the eSyn-OPC vicinity. Given that each Visium capture spot (55 μm diameter) encompasses multiple cells, we reasoned that the transcriptomic composition of eSyn-OPC spots would reflect the identity of spatially proximal cell types [33]. We performed differential expression analysis between *MT1E*^+^ spots (containing eSyn-OPCs) and *PDGFR*α^+^/*MT1E*^-^ spots (containing conventional OPCs but not eSyn-OPCs). This comparison demonstrated that spots containing eSyn-OPCs preferentially expressed markers of neural progenitors (*EOMES*) and neural stem cells (*SOX2, ID4*), suggesting a physical proximity to these lineages (**Fig. 4H**). Importantly, this spatial association with neural progenitors/stem cells persisted at the later developmental stage of PCW 19 (**Suppl Fig. 4G**).

To validate these spatial interactions, we performed triple immunohistochemistry on PCW 22 tissue. This revealed eSyn-OPCs (GRB14^+^/PDGFRα^+^) residing in immediate adjacency to SOX2^+^ neural stem cells (**Fig. 4I**), suggesting a potential direct, or proximal communication.

### eSyn-OPCs are oligodendrocyte lineage cells capable of differentiation

To further validate the cellular lineage of eSyn-OPCs, we searched the scRNA-seq data for unique cell surface markers for eSyn-OPCs. Out of 1,777 proteins annotated as “cell membrane” or “plasma membrane” proteins in human UniProt database, we identified GRB14 as a unique marker. It is both highly expressed as well as enriched specifically in eSyn-OPCs (**Fig. 5A**, **Suppl Fig. 4B-C**). We optimized an antibody-based immunopanning protocol to isolate eSyn-OPCs based on double positivity for GRB14 and PDGFRα (**Fig. 5B**). The isolated cells confirmed GRB14 and PDGFRα immunoreactivity along with OLIG2, another OPC marker, upon plating on coverslips (**Fig. 5C**). Once they were induced to differentiate with differentiation medium lacking growth factors, eSyn-OPCs differentiated to O4^+^/MBP^+^/PLP1^+^ mature oligodendrocytes (**Fig. 5D**). These data demonstrate that eSyn-OPCs are indeed oligodendrocyte lineage cells that are capable of differentiating to mature oligodendrocytes when directed *in vitro*.

**Figure 5.**
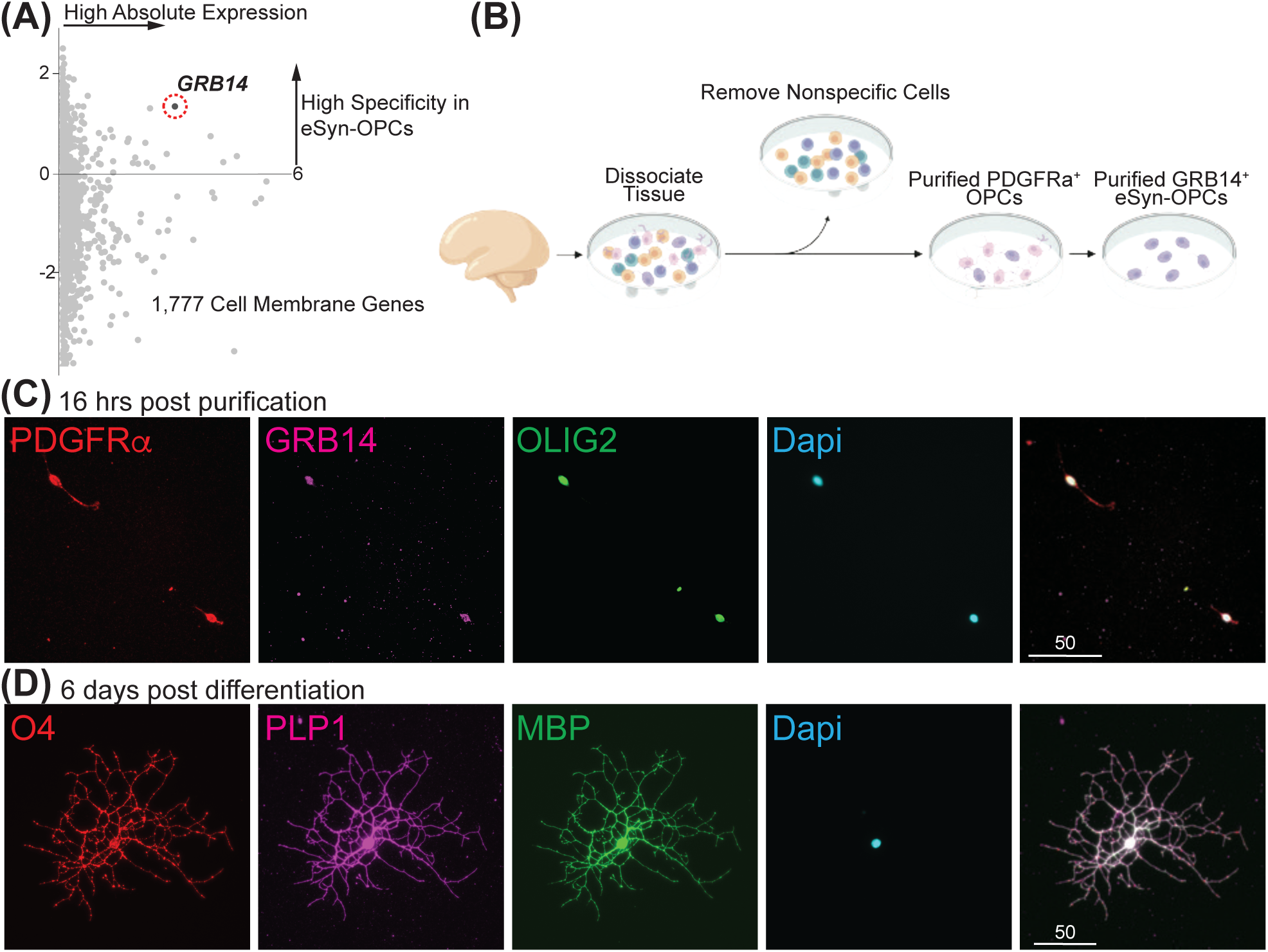
Validation of eSyn-OPC cellular lineage. **(A)** Enrichment in cell surface marker genes in eSyn-OPCs compared to non-Syn-OPCs. GRB14 was identified as a unique marker. **(B)** A schematic illustrating the isolation of eSyn-OPCs based on double positivity for GRB14 and PDGFRα antibodies. **(C)** The isolated eSyn-OPCs are GRB14^+^/PDGFRα^+^/OLIG2^+^. Scale bar unit = um. **(D)** When isolated eSyn-OPCs were induced to differentiate with differentiation medium lacking growth factors, eSyn-OPCs differentiated into O4^+^/MBP^+^/PLP1^+^ mature oligodendrocytes. Scale bar unit = um.

## DISCUSSION

Our observations highlight that during the second trimester of human development, distinct clusters of OPCs are found in the cortex. One cluster, eSyn-OPC, emerges sometime between PCW 12 and 15 (**Fig. 4B-C, E-F, Suppl Fig. 4D-F**). Although the proportion of eSyn-OPCs vary across cortical layers, VZ, iSVZ and oSVZ are the most enriched. Based on the enrichment in VZ in younger tissue (PCW 15) compared to the enrichment in oSVZ in older tissue (PCW 22) (**Fig. 4B-C**), it is possible that eSyn-OPCs are generated in VZ and migrate towards oSVZ. In all tissues we examined (N = 7), negligible amounts of eSyn-OPCs were observed in cortical layers above oSVZ (**Fig. 4B-C, E-F, Suppl Fig. 4D-F**), although PDGFRα^+^ OPCs are abundant (**Fig. 4B, G, Suppl Fig. 4D-F**). It is possible that the ventral layers provide an optimal environment for the syn-OPCs, as the timing of the clear appearance of oSVZ at PCW 13 and onward [19] correlates with the appearance of eSyn-OPC in our samples. However, it is also important to note that due to the lack of regional information in the tissues that we obtain, it is possible that eSyn-OPCs are region-specific, and we simply obtained cortical regions at below PCW 15 that the eSyn-OPCs do not exist.

We identified an adult counterpart, aSyn-OPCs that show strong evidence of roles in synaptic regulation, implicating a possibility that eSyn-OPCs persist into adulthood as aSyn-OPCs. On the other hand, despite the similarities in pathway signatures between eSyn-OPCs and aSyn-OPCs (**Fig. 1H, 2F, Fig. 3B**), the overlap of the DEGs in these clusters was limited to 5.52% (**Fig. 3A**), suggesting functional similarity needs to be tested.

Accumulating reports indicate that mouse OPCs in various brain areas such as motor cortex, corpus callosum, thalamus, and somatosensory cortex receive monosynaptic connection [34], and that OPCs sense excitatory and inhibitory neurotransmitters with functional receptors via neuron-OPC synapse [35, 36]. Expression of various neurotransmitter receptor subunits have been reported in rodents (reviewed in [37, 38]) and human [39], although subunit types may differ between species. Neuronal activity-dependent OPC proliferation and differentiation is well documented in mouse [40–42] and zebrafish [43–45] during postnatal myelination. We therefore used SynGO [32], the curated synapse-specific gene annotation, to ask whether human eSyn-OPCs and aSyn-OPCs may possess similar characteristics. We found that eSyn-OPCs highly expressed group of genes that encode for postsynaptic proteins (**Fig. 3C**) especially neurotransmitter receptor subunits for glutamate, GABA, and acetylcholine (**Fig. 3D-F**), suggesting eSyn-OPCs may be able to sense these neurotransmitters. If so, it implies OPC-neuron communication at an early stage of human development, in contrast to the reported findings in animal models [46].

Other groups have reported that, in developing and adult mouse visual cortex, a subset of OPCs engage in axon pruning by engulfing synapses [47, 48]. Similar report was made in zebrafish optic ganglion where ablation of OPCs in this region impaired neurite pruning and remodeling [49]. We therefore tested whether the genes related to phagocytosis highly expressed by mouse OPCs were also expressed by eSyn-OPCs, but they did not express any of the 38 genes [47] at noticeable level, implying eSyn-OPCs may not have phagocytic capacity but rather use other mechanisms to impose synaptic regulation. In contrast, 2 out of 38 genes related to phagocytosis were expressed in aSyn-OPCs (data not shown). Taken together, the current study reports clearly defined OPC populations that are evident during human development as well as adulthood, and that the Syn-OPC population may potentially play a role in synaptic signaling from early developmental stages.

## METHODS

### Fetal brain tissues

Specimens from fetal (PCW 8 to 22) human brain tissues were obtained from the Birth Defects Research Laboratory at the University of Washington with ethics board approval and maternal written consent. This study was performed in accordance with ethical and legal guidelines of the Rutgers University’s institutional review board.

### Isolation of OPCs

Fetal brain tissues were dissociated with papain for 10min. The single cell suspension was kept in PDGF medium containing DMEM/F12 (Thermofisher, 10565018), non-essential amino acids (Thermofisher, 11140050), N2 supplement (Thermofisher, 17502-048), B27 supplement (Thermofisher, 17504044), Penicillin-Streptomycin (Thermofisher, 15-070-063), Insulin (Sigma, I9278-5ML, final conc 25ug/ml), PDGF-AA (Peprotech, 100-13A, final concentration 10ng/ml), IGF (Peprotech, 100-11, final concentration 10ng/ml), NT3 (Peprotech, 450-03, final concentration 1ng/ml), Biotin (Sigma, B4639, final concentration 100ng/ml), cAMP (Sigma, D0260-5MG, final concentration 1uM), and T3 (Sigma, T6397, final concentration 40ng/ml). Antibody-coated dishes were prepared by first incubating 200ug of appropriate secondary antibody (Jackson Immunoresearch) in Ph 9.5 Tris buffer overnight at 4C. The plate was rinsed in PBS 3 times, then 40ug primary antibody was incubated in PBS containing 0.1% BSA for at least 1hr. The plate was rinsed in PBS 3 times before dissociated cells were added. Dissociated cells were passed onto a dish coated with mouse IgG secondary antibody only (Jackson Immunoresearch, 115-005-003) for 2hr at 37C to avoid non-specific binding, then floating cells were transferred to a dish coated with PDGFRα antibody (BD Biosciences, 556001) for 35min at 37C, to isolate PDGFRα^+^ OPCs. At the end of PDGFRα binding, plates were washed to remove non-adherent cells, then adherent cells were detached with Trypsin, and collected by centrifugation. To isolate eSyn-OPCs, PDGFRα^+^ OPCs were passed on to a dish coated with GRB14 antibody (Proteintech, 15298-1-AP) for 20min at 37C. Plates were washed to remove non-adherent cells, then adherent cells were detached with Trypsin, collected by centrifugation, and plated on to poly-L-ornithine/laminin-coated coverslips.

### Differentiation of isolated eSyn-OPCs

Isolated and plated eSyn-OPCs were differentiated by changing the PDGF medium to glia medium containing DMEM/F12 (Thermofisher, 10565018), non-essential amino acids (Thermofisher, 11140050), N2 supplement (Thermofisher, 17502-048), B27 supplement (Thermofisher, 17504044), Penicillin-Streptomycin (Thermofisher, 15-070-063), Insulin (Sigma, I9278-5ML, final conc 25ug/ml), Biotin (Sigma, B4639, final concentration 100ng/ml), cAMP (Sigma, D0260-5MG, final concentration 1uM), T3 (Sigma, T6397, final concentration 40ng/ml), and ascorbic acid (Sigma, A4544, final concentration 20ug/ml) for 6 days, with 50% medium change every other day.

### Immunohistochemistry

Tissues or cells were fixed in 4% paraformaldehyde, and immunocytochemistry was performed with antibodies to PDGFRα (BD Biosciences, 556001), HAPLN1 (LS Bio, LS-C334832-20), IL1RAP (Proteintech, 21609-1-AP), SPOCK2 (Proteintech, 11725-1-AP), GRB14 (Proteintech, 15298-1-AP), and MT1E (Millipore Sigma, HPA041152-25UL) overnight at 4C. Secondary antibodies of appropriate species were purchased from ThermoFisher.

### Quantitative Analysis

Fluorescent images were captured by Leica DMi8 microscope equipped with a sCMOS camera. Dapi was used to delineate cortical layers. 20X images taken from 4 technical replicates in each sample were manually analyzed for the proportion of PDGFRα^+^/HAPLN1^+^ eSyn-OPCs relative to all PDGFRα^+^ OPCs in randomly selected fields. At least 300 PDGFRα^+^ cells were used for quantification in each sample.

### scRNA-seq (embryonic OPCs)

Single-cell suspensions of isolated OPCs from a PCW 17 cortical specimen were processed using the Chromium Next GEM Single Cell 3’ v2 Reagent Kits (10x Genomics) following the manufacturer’s instructions. Briefly, cells were resuspended at a concentration of 440 cells/μl, and ~100,000 cells were loaded onto a Chromium Controller to generate single-cell Gel Bead-in-emulsion (GEMs). cDNA amplification and library construction were performed according to standard protocols, and library quality was assessed using an Agilent TapeStation. Sequencing was conducted on an Illumina HiSeq 4000 platform to achieve a targeted depth of 100,000 reads per cell. Raw FASTQ files were aligned to the human reference genome using the Cell Ranger *count* pipeline (v3.0.2) with default parameters. The raw FASTQ files are available via the NCBI Gene Expression Omnibus (GEO) under sample ID: GSM6598840. Please note that this sample is annotated as 19 week gestational age in the database, corresponding to PCW 17.

### snRNA-seq (adult OPCs)

To investigate the conservation of OPC subtypes in adulthood, we analyzed publicly available single-nucleus RNA-seq datasets from adult human donors with no known history of neuropsychiatric or neurological conditions. Raw FASTQ files were downloaded from the Neuroscience Multi-omics (NeMO) Archive (RRID:SCR_016152) and aligned to the GRCh38-2020-A reference genome using the Cell Ranger count pipeline (v7.0.1) with default parameters. Individual output files were integrated separately for the 29-year-old male and 42-year-old male donors using Cell Ranger aggr with the “--normalize=none” setting. Quality control was performed on the Schirmer et al. dataset (**Suppl Fig. 3**) based on the Fraction Reads in Cell matrix to identify and exclude outlier libraries, defined as values falling below 65%. Consequently, 16 out of 21 libraries passed the QC check and were retained for secondary analysis. The OPC supercluster was subsequently identified based on the expression of *PDGFRA* and *OLIG2*. These cells were subsetted and re-clustered using the reclustering workflow within the Loupe Browser’s Reanalyze panel. Subclusters were annotated based on marker gene expression and Gene Ontology (GO) enrichment analysis performed using Metascape. The custom Python script used for extracting quality control metrics from Cell Ranger outputs is available at https://github.com/kosaku-san.

### Spatial transcriptomics

Spatial transcriptomic analysis was performed using the Visium Spatial Platform (10x Genomics) on fresh-frozen cortical tissues from PCW 8, 15, and 19. Tissue sections (10um thickness) were placed onto the active capture areas of Visium slides. The sections were fixed with methanol, stained with hematoxylin and eosin, and imaged using a brightfield microscope to capture morphological context. Tissue permeabilization was optimized to 18 minutes. On-slide cDNA synthesis, second-strand synthesis, and denaturation were performed according to the manufacturer’s instructions. Libraries were constructed following the standard protocol and sequenced on an Illumina NovaSeq 6000 platform. Raw FASTQ files and corresponding images were processed using the Space Ranger count pipeline (v1.3.1) with default parameters. Reads were aligned to the GRCh38-2020-A reference genome. Output files from individual runs were combined using Space Ranger aggr (v2.0.0) with the “--normalize=mapped” setting. The raw FASTQ files for each dataset are accessible via the NCBI GEO under accession number GSE213306, with the corresponding sample IDs: GSM6578640 (PCW 8), GSM6578643 (PCW 15), and GSM6578645(PCW 19). Please note that in the GEO database, sample ages are annotated using gestational week rather than post-conceptional week (PCW).

### Monte Carlo Simulation

To assess the statistical significance of the overlap between embryonic eSyn-OPC and adult aSyn-OPC markers, we performed a Monte Carlo simulation [30]. The observed overlap of the top 800 marker genes was compared against a null distribution generated by randomly sampling gene sets of equal size from a background of expressed cortical genes. This background gene pool was defined using The Human Protein Atlas, restricted to genes with expression levels of 1 transcripts per million (TPM). The source code for this analysis is available at https://github.com/kosaku-san.

## ACKNOWLEDGEMENTS

Research reported in this publication was supported by the National Institute of Neurological Disorders and Stroke of the National Institutes of Health (NIH) under award number 5R21NS132013 (to H.N.), National Multiple Sclerosis Society award number TA-1704-27504 (to H.N.), and Rutgers University’s Busch Biomedical Grant (to H.N.). K.S. is supported by the NIH under award numbers DK110426, DK110063, and DK128839, and the American Federation for Aging Research (AFAR) under grant numbers 21139 and 22059. We also acknowledge the DERC (NIH P30 DK020541) as well as NORC (P30 DK026687) grants for single-cell genomics. H.N. acknowledges the Birth Defects Research Laboratory (BDRL) for human brain tissue collection. BDRL is funded by NIH under NICHD Grant # R24HD000836 to I.A.G. Schematic illustrations included in Figure 5B were created using BioRender by H.N. (agreement number LO292MNNLL). We thank David M. Reynolds at the Genomics Core at Albert Einstein College of Medicine for assistance in obtaining spatial transcriptomic data from human brain tissues. We also thank Peter Lönnerberg and Kimberly Siletti at Karolinska Institute, Sweden, for technical assistance in data analysis.

---

**Supplementary Figure 1.**
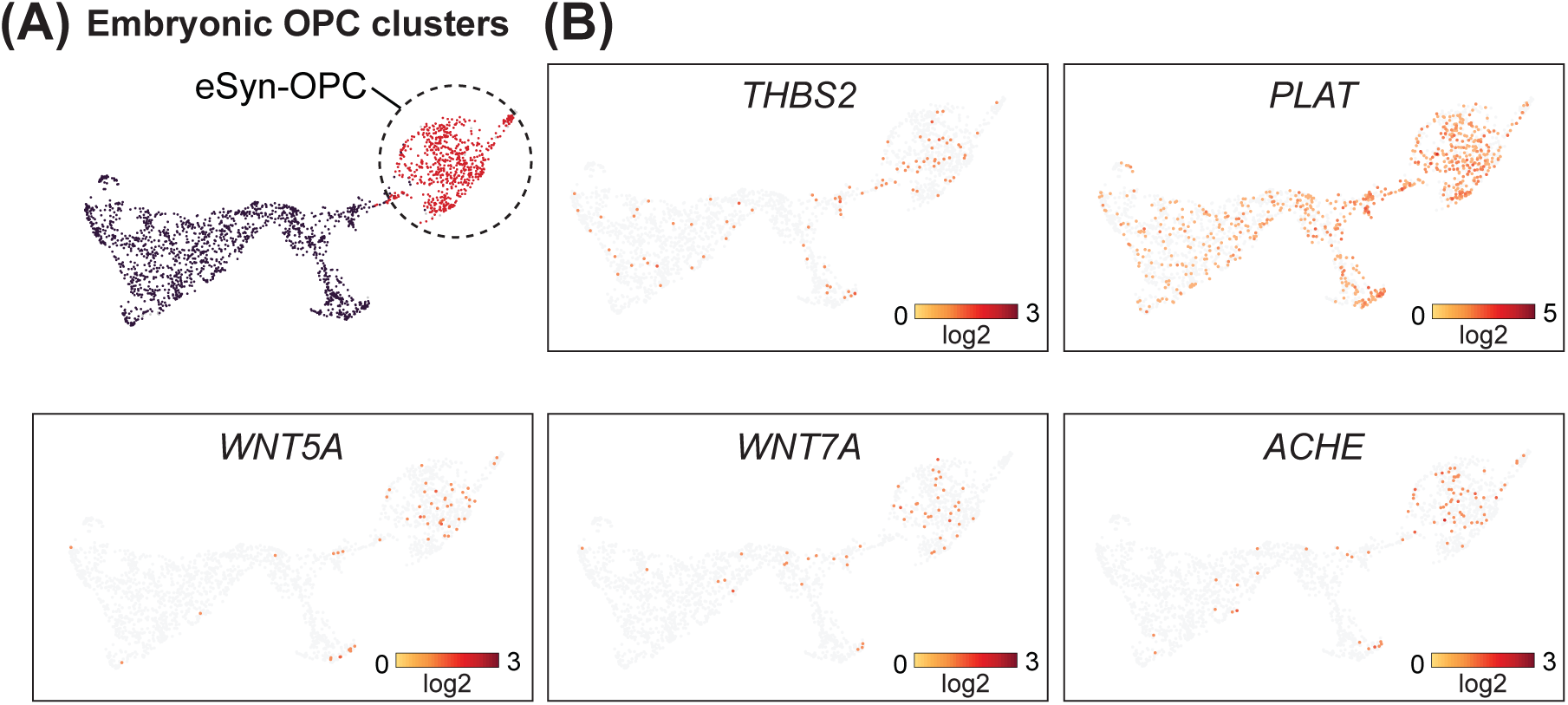
eSyn-OPCs express known synaptic development genes. **(A)** (Top left) Graph clustering of eSyn-OPCs in UMAP space from the PCW 17 brain, highlighted in red. **(B)** eSyn-OPCs express known synaptic development genes, including *THBS2, PLAT, WNT5A, WNT7A,* and *ACHE*.

**Supplementary Figure 2.**
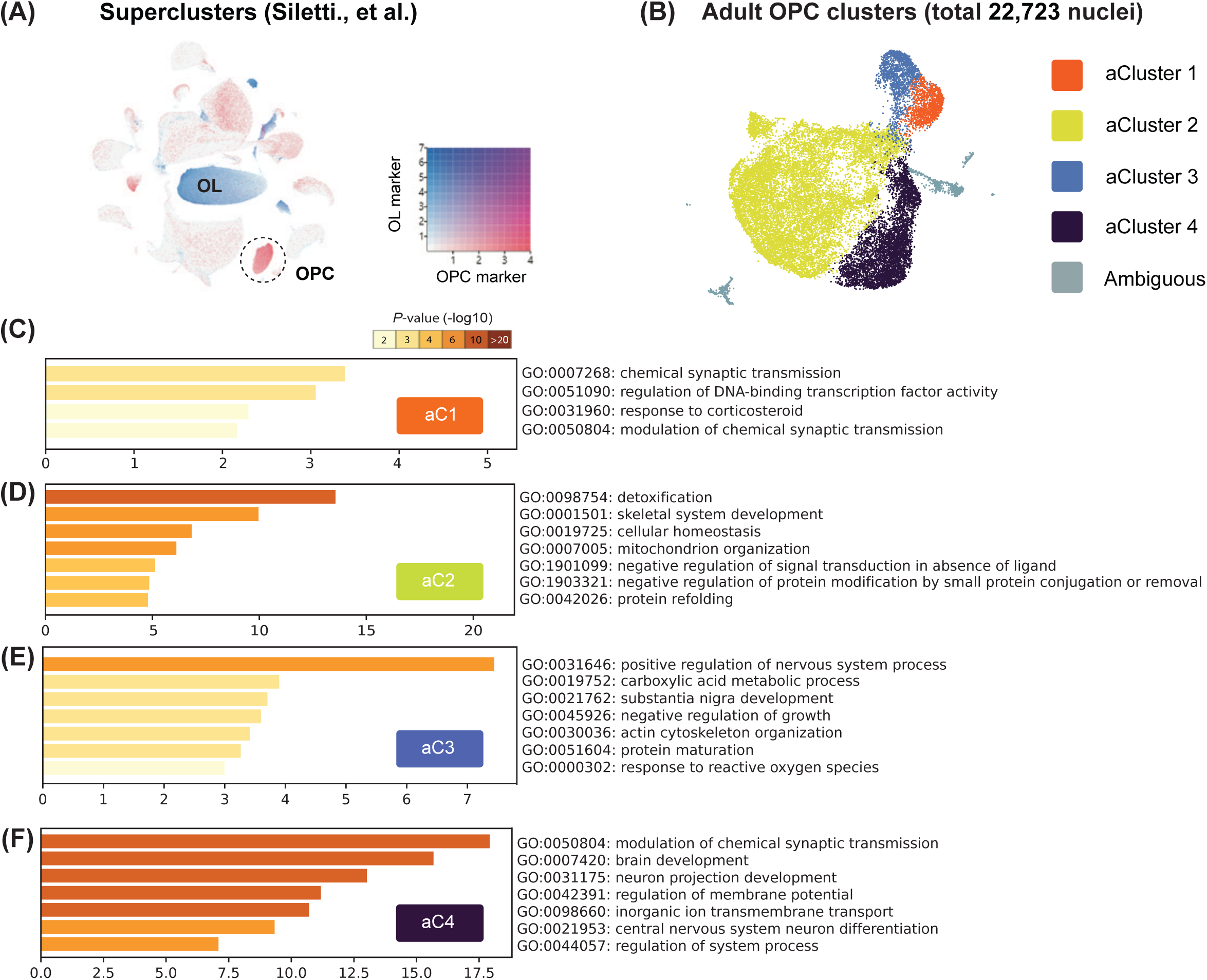
Evidence of aSyn-OPCs in another adult human brain. **(A)** Identification of OPCs in the UMAP plot of brain cells obtained from a healthy 42-year-old male. OPCs were defined by co-expression of marker genes *PDGFRA* and *OLIG2*, and mature oligodendrocytes were defined by co-expression of marker genes *MAG*, *MOG*, and *MBP*. **(B)** Re-clustering of the OPC cluster found in (A) based on differentially expressed genes. **(C-F)** Top pathways identified by GO analysis of differentially expressed genes in each OPC cluster from (B). <u>A</u>dult Cluster 4 (aC4) was identified as aSyn-OPCs.

**Supplementary Figure 3.**
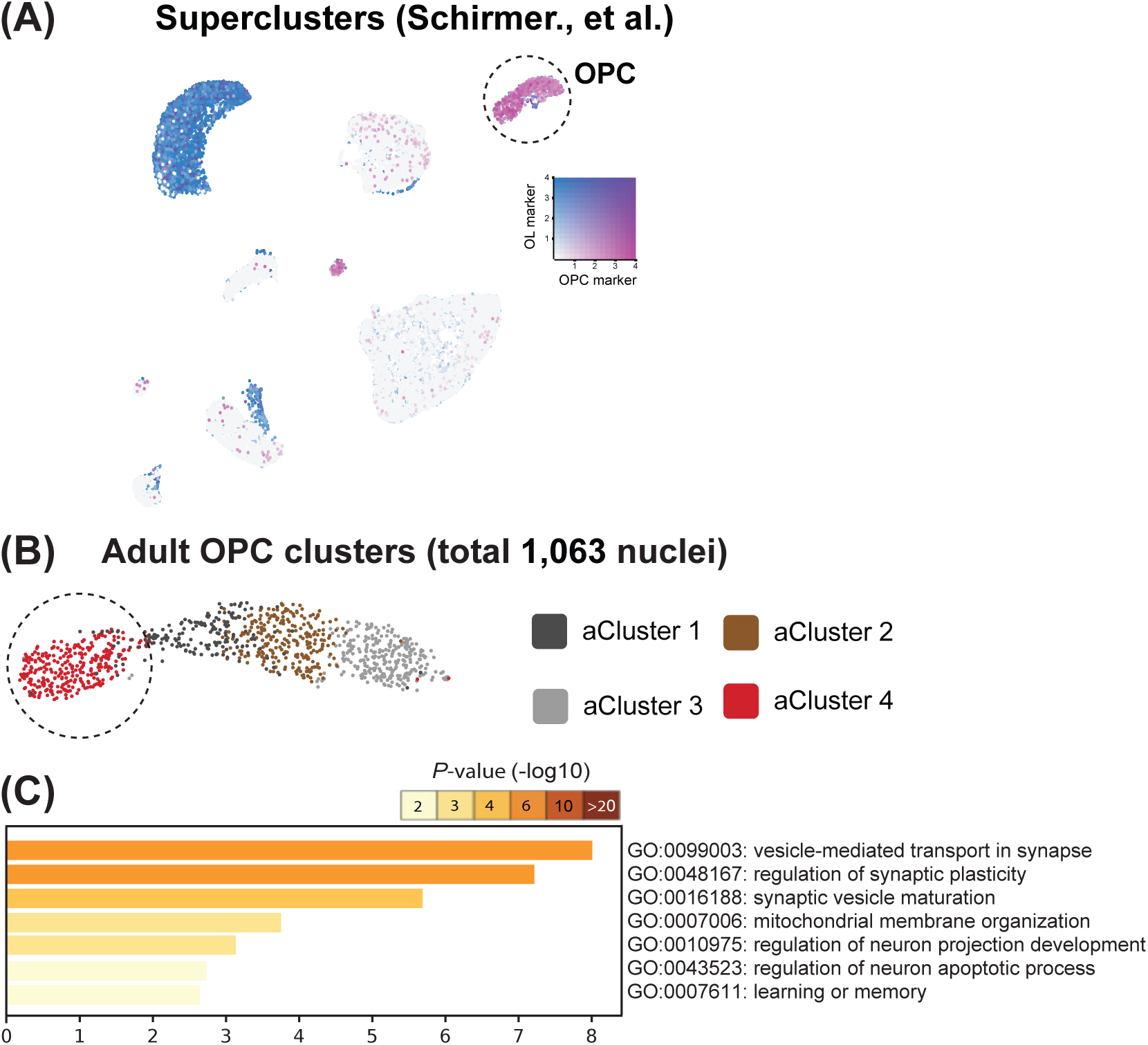
Evidence for synaptic OPCs across adult age groups. **(A)** An independent dataset from Schirmer et al. was used to identify OPCs from the single-nucleus RNA sequencing of cortical brain cells obtained from 6 healthy individuals of various ages (34 - 68 years old). OPCs were defined by co-expression of marker genes *PDGFRA* and *OLIG2*, and mature oligodendrocytes were defined by co-expression of marker genes *MAG*, *MOG*, and *MBP*. **(B)** Re-clustering of the OPC cluster found in (A) based on differentially expressed genes. **(C)** Top seven pathways identified by GO analysis of differentially expressed genes in <u>A</u>dult Cluster 4 (aC4), identified as aSyn-OPCs.

**Supplementary Figure 4.**
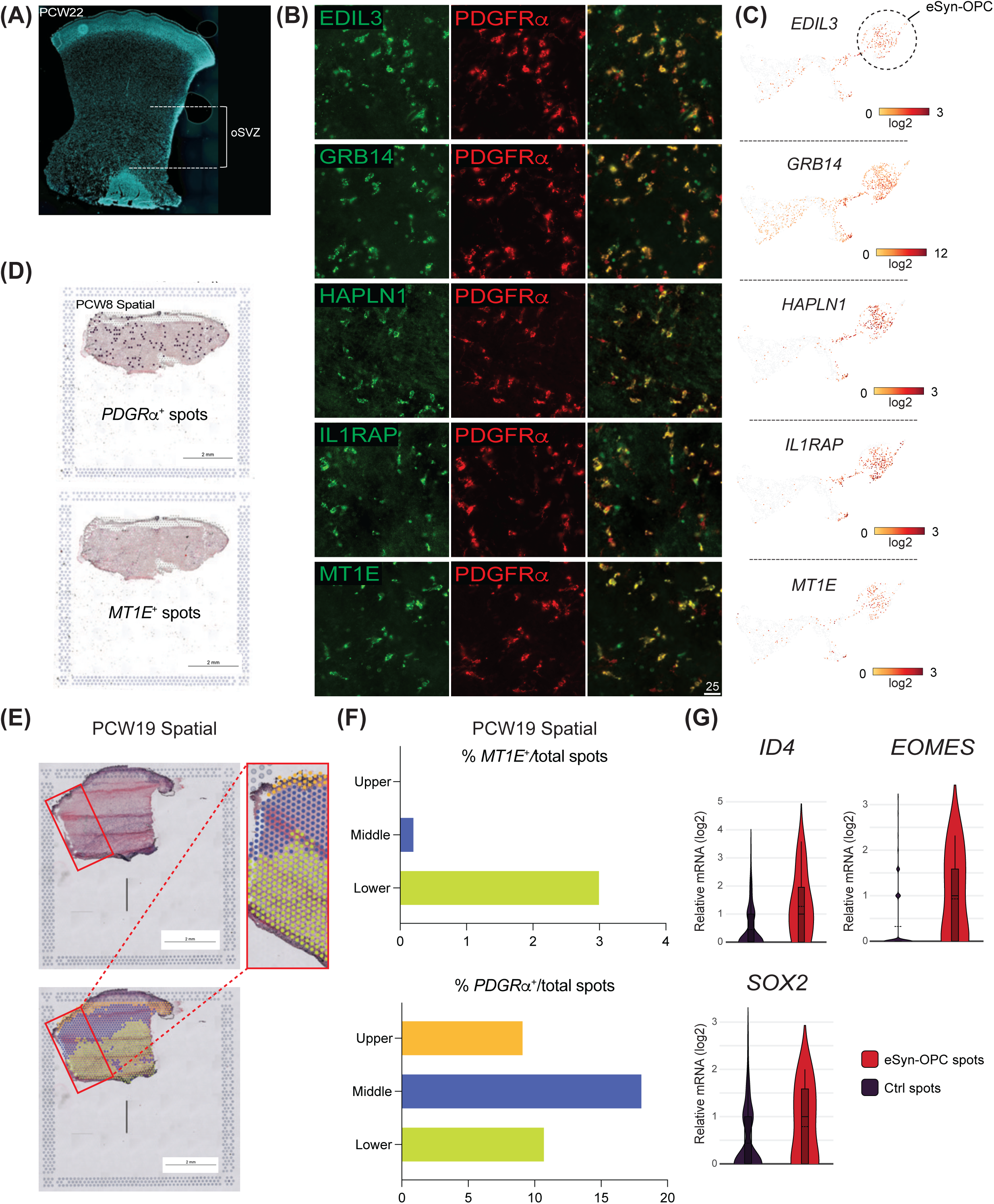
eSyn-OPC markers that are validated by staining and gene expression. **(A)** A representative image of a cortical section obtained from a PCW 22 sample. The location of the oSVZ region is indicated. **(B)** Immunohistochemistry with candidate eSyn-OPC markers. Images were taken from adjascent sections in the oSVZ region indicated in (A). Scale bar unit = um. **(C)** Corresponding mRNA expression levels of candidate markers used in the immunohistochemistry shown in (B). **(D)** A spatial transcriptomic sample from PCW 8 tissue. Negligible *MT1E*^+^ spots (8 spot; 0.7%) were found despite the abundant PDGFRα^+^ spots. **(E)** A spatial transcriptomic sample from PCW 19 tissue. **(F)** *MT1E*^+^ spots were enriched in lower cortical layers whereas PDGFRα^+^ spots were found across cortical layers. **(G)** The mRNA levels of neural progenitor (*EOMES*) and neural stem cell (*SOX2, ID4*) markers in the *MT1E*^+^ spots (containing eSyn-OPCs) compared to *PDGFR*α^+^/*MT1E*^-^ spots (containing OPCs but not eSyn-OPCs) in PCW 19 sample.

